# DiscoTope-3.0 - Improved B-cell epitope prediction using AlphaFold2 modeling and inverse folding latent representations

**DOI:** 10.1101/2023.02.05.527174

**Authors:** Magnus Haraldson Høie, Frederik Steensgaard Gade, Julie Maria Johansen, Charlotte Würtzen, Ole Winther, Morten Nielsen, Paolo Marcatili

## Abstract

Accurate computational identification of B-cell epitopes is crucial for the development of vaccines, therapies, and diagnostic tools. However, current structure-based prediction methods face limitations due to the dependency on experimentally solved structures. Here, we introduce DiscoTope-3.0, a markedly improved B-cell epitope prediction tool that innovatively employs inverse folding structure representations and a positive-unlabelled learning strategy, and is explicitly adapted for both solved and predicted structures. Our tool demonstrates a considerable improvement in performance over existing methods, accurately predicting linear and conformational epitopes across multiple independent datasets. Most notably, DiscoTope-3.0 maintains high predictive performance across solved, relaxed and predicted structures, alleviating the need for experimental validation and extending the general applicability of accurate B-cell epitope prediction by more than 3 orders of magnitude. DiscoTope-3.0 is made widely accessible on two web servers, processing over 100 structures per submission, and as a downloadable package. In addition, the servers interface with RCSB and AlphaFoldDB, facilitating large-scale prediction across over 200 million cataloged proteins. DiscoTope-3.0 is available at: https://services.healthtech.dtu.dk/service.php?DiscoTope-3.0

## 1 Introduction

A key mechanism in humoral immunity is the precise binding of B-cell receptors and antibodies to their molecular targets, named antigens. The antigen regions that are involved in the binding are known as B-cell epitopes. B-cell epitopes are found on the surface of antigens, and in the case of proteins they can be classified as linear if the epitope residues are sequentially arranged along the antigen sequence, or discontinuous if they are only proximal in the antigen tertiary structure, but not in the primary structure. Identification of B-cell epitopes has large biotechnological applications, including rational development of vaccines and immunotherapeutics. However, experimental mapping of epitopes remains expensive and resource intensive. Computational tools for B-cell epitope prediction offer a viable and large-scale alternative to experiments.

However, prediction of B-cell epitopes remains a challenging problem (Galanis et al., 2019; Sun et al., 2019). Historically, in-silico prediction methods have been either antigen sequence or structure-based. Sequence-based methods such as BepiPred-2.0 (Jespersen et al., 2017) are attractive given the high availability of protein sequences. BepiPred-2.0 utilizes a random forest trained on structural features predicted from the antigen sequence, but has limited accuracy and struggles to predict conformational or non-linear epitopes (Klausen et al., 2019). In a recent work, BepiPred-3.0 (Clifford et al., 2022) further improves the method, demonstrating large gains by exploiting sequence representations from the protein language model ESM-2 (Lin et al., 2022). It was shown to outperform previous sequence based tools, including Seppa-3.0 (Zhou et al., 2019), ElliPro (Ponomarenko et al., 2008), BeTop (Zhao et al., 2012), EPSVR, EPMeta (Liang et al., 2010), and EPSVR / EPMeta (Liang et al., 2010).

Structure-based methods should benefit from having direct access to the antigen tertiary structure, and in particular, its surface topology. DiscoTope-2.0 (Kringelum et al., 2012) was published in 2012, and it estimates epitope propensity from the local geometry of each residue, taking into consideration both its solvent accessibility and the direction of its side chain. Older structure-based methods like DiscoTope-2.0 and the newer epitope3D (da Silva et al., 2021) are still outperformed by the sequence-based BepiPred-3.0 (Clifford et al., 2022). However novel methods such as the inverse-folding based SEMA (Shashkova et al., 2022) and the geometric deep-learning network ScanNet (Tubiana et al., 2022b) have shown promising advances. Recently, ScanNet demonstrated improved performance by explicitly considering geometric details at both the resolution of individual atoms and amino-acids. However, while structure-based prediction tools may demonstrate improved performance, they are limited by the availability of antigen structures.

Data scarcity affects the accuracy of prediction tools in different ways. Firstly, they constrain the amount of data on which such tools can be trained. As of January 2023, less than 5500 antibody structures in complex with an antigen are available in the antibody-specific structural database SabDab (Dunbar et al., 2013). After filtering this dataset for redundancy, one may be left with less than 1500 structures for training, which limits the complexity of the models that can be reliably trained without incurring in overfitting (Clifford et al., 2022).

Secondly, the available data is a biased sampling of the possible antibody-antigen complexes. We find that most antigens are found only once in the dataset, while others, likely due to medical or biological interest, have been resolved in complex with as many as 43 (Dunbar et al., 2013) different antibodies. This means that one cannot confidently annotate negative residues; they might be part of antibody-antigen complexes yet to be solved.

Lastly, undersampling of epitopes will also result in an imprecise assessment of the tools’ accuracy; predicted epitopes that appear as false positives may just be in antibody bound regions yet to be identified. The last two points (bias and undersampling) are typical of a class of problems known as Positive-Unlabeled (PU) learning. In this scenario, we are only confident of positive epitopes, while all remaining (surface) residues should be treated as unlabeled. Several approaches have been proposed for increasing the accuracy of B-cell epitope prediction methods and their estimated metrics in such cases (da Silva et al., 2021; Ren et al., 2015; Li et al., 2021). A simple yet effective strategy is to train ensemble predictors based on bootstrapping of samples in the Unlabelled class (Mordelet and Vert, 2014), also known as PU bagging, which is the approach that we propose in this work.

With recent advances in protein structure prediction, AlphaFold2 (Jumper et al., 2021) has enabled accurate prediction of protein structures directly from sequences. Currently, over 200 million pre-computed structures are available in AlphaFold DB (Varadi et al., 2021), covering every currently cataloged protein in UniProt (Consortium, 2022). The three-dimensional coordinates of the proteins, together with the local quality reported as pLDDT scores, are readily accessible from the database.

To truly harness the remarkable progress in generating accurate structural models, we must develop robust and informative numerical representations of both predicted and resolved structures. This is especially crucial for deep-learning methods, which thrive on such tasks. The ESM-IF1 inverse folding model is an equivariant graph neural network pre-trained to recover native protein sequences from protein backbones structures (Ca, C and N atoms). The structure-based representations which may be extracted from this model have been shown to outperform sequence-based representations on tasks such as predicting binding affinity and change in protein stability (Hsu et al., 2022). Crucially, ESM-IF1 is explicitly trained on both solved and AlphaFold predicted structures, enabling large-scale application of its representations even when solved structures are unavailable.

In this work, we train DiscoTope-3.0, a structure-based B-cell epitope prediction tool exploiting inverse folding representations generated from either AlphaFold predicted or solved structures. DiscoTope-3.0 is explicitly trained on both predicted and solved antigen structures using a positive-unlabelled learning ensemble approach, enabling large-scale prediction of epitopes even when solved structures are unavailable. We compare it’s performance versus previous tools and the impact in performance when using predicted structures versus solved structures, in both cases showing unprecedented accuracy. DiscoTope-3.0 is implemented as a web server and downloadable package interfacing with both RCSB and AlphaFoldDB.

## 2 Results

The positive-unlabelled ensemble training strategy for DiscoTope-3.0 is shown in Figure 1. First, epitopes from solved antibody-antigen complexes are mapped onto the antigen sequences (1). Using sequences as input, antigen structures are predicted using AlphaFold2 (2). Next, per-residue structural representations, for both solved and predicted structures, are extracted using the ESM-IF1 protein inverse folding model (3 and 4). During training, random subsets of epitopes and unlabelled residues are sampled across the dataset (5), before finally training an ensemble of XGBoost models on the individual data subsets (6). The final DiscoTope-3.0 score is given as the average score from the ensemble models (7). More details on the training procedure are available below and in the Methods section.

**Fig. 1.**
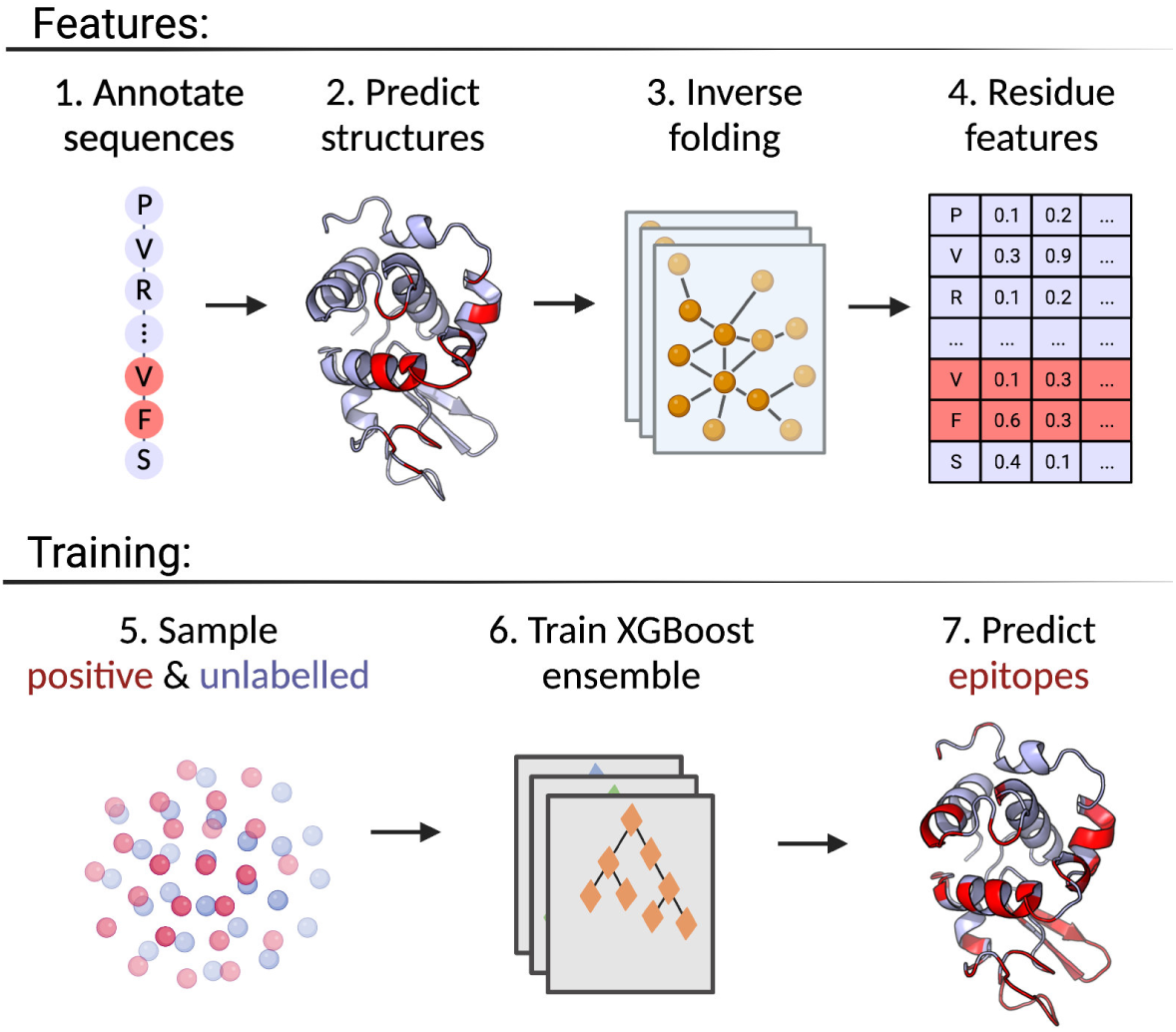
Overview of the DiscoTope-3.0 method. Created with BioRender.com

Here, we present a quick overview of the dataset and feature pre-processing procedure. DiscoTope-3.0 training and validation is based on the BepiPred-3.0 dataset of 582 antibody-antigens complexes, covering a total of 1466 antigen chains IEDB (Vita et al., 2018). Epitopes are defined as the set of residues within 4 Å of any antibody heavy atom (see Methods). The training and hyperparameter tuning is based on 2 different datasets: Training and Validation, while evaluation is performed on the Validation and external test sets. The external test set consists of 24 antigens collected from SAbDab (Dunbar et al., 2013) and PDB (Berman et al., 2000) on October 20, 2022. These antigens share at most 20 % similarity to both our own, BepiPred and ScanNet’s training datasets (see Methods).

In addition to using experimentally solved antigens for training, all individual antigen chains were predicted using AlphaFold2. Both the solved and predicted chains were then embedded with ESM-IF1. Further we extract for each residue its relative surface accessibility (RSA), AlphaFold local quality score (pLDDT) as well as the antigen length and a one-hot encoding for the antigen sequence (see Methods and Supplementary table 1). These structural features (or subsets) were used to train an ensemble of XGBoost models and the ensemble average is used as the final prediction score.

We chose to use XGBoost for our architecture due to their robustness to outliers and noise, minimal need to adjust model hyperparameters (Chen and Guestrin, 2016), and enabling combination of multiple ”weak learners” in our PU (positive and unlabelled) learning ensemble (Zhao et al., 2022; Claesen et al., 2015), to produce a robust final prediction.

Structure-based representations have been shown to be a powerful representation in different downstream tasks. To see if this is also the case for B-cell epitope prediction, we evaluated the results obtained using different feature encoding schemes on our validation set of AlphaFold structures (for details on this dataset refer to Methods). First, we assess whether training a single XGBoost model using structure representations from predicted structures outperforms a similar model based on the sequence representations from ESM-2 (Figure 2). Here, we observe a marginal but consistent epitope prediction performance using the structure (AUC-ROC 0.767 *±* 0.003) vs sequence representations (AUC-ROC 0.751 *±* 0.003) (*p <* 0.0001).

**Fig. 2.**
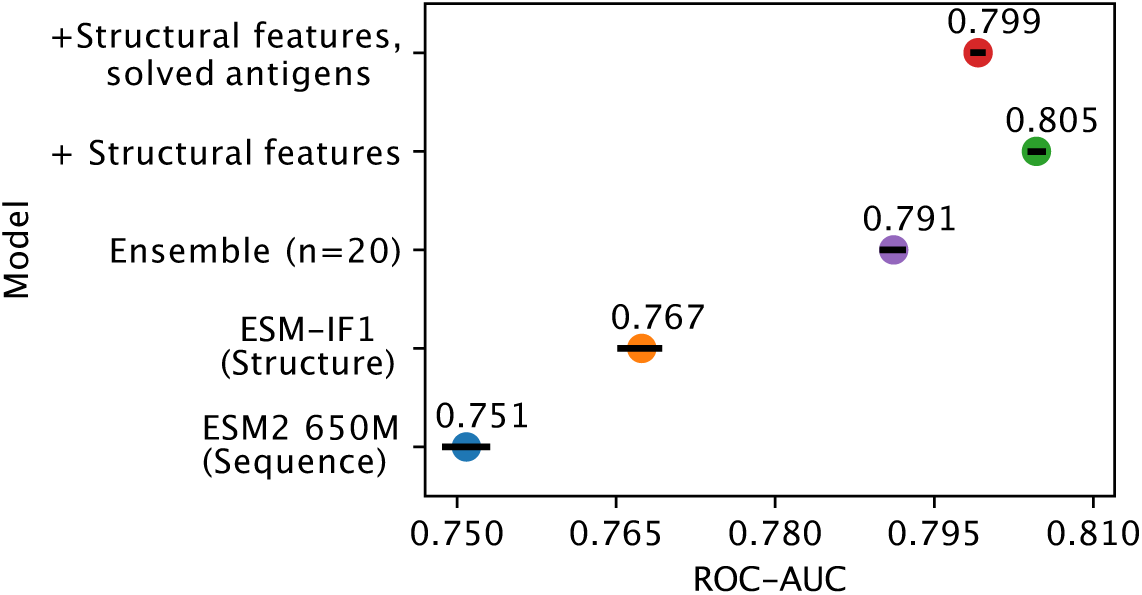
Effects of inverse folding and bagging. Ablation results on the validation set of AlphaFold predicted structures, with less than 50 % sequence similarity to the training set. The plot reports the AUC-ROC for a single XGboost model trained on representations based on ESM-2 650M parameter (blue) and on ESM-IF1 (orange), for an ensemble of 20 XGboost models based on bootstraps of ESM-IF1 representations (purple), for models where additional structural features are included (see methods) tested on both AlphaFold models (green) and on the corresponding solved structures (red) (see Methods). Error bars indicate 95 % confidence interval.

As explained in the introduction, the B-cell epitope prediction problem can be categorized in the broad class of PU training. Incorrectly labeled negative examples can negatively affect the training, by introducing frustration in the learning process (Dietterich, 2000). We can observe that, by using an ensemble learning strategy with a dataset bagging approach based on previous works (Huang et al., 2009; Elkan and Noto, 2008; Dietterich, 2000) (see Methods), we can further improve performance (AUC-ROC 0.791 *±* 0.001) and generalization.

### 2.1 Effect of using predicted versus solved structures

One of the risks in training models on either exclusively solved or AlphaFold structures is that the methods might over-specialize to one source and perform worse on the other, or even be affected by data leakage. For example, a model may overfit on conformational changes present in the side chains of epitope residues in solved antibody-antigen complexes.

By training on both predicted and solved structures, we obtain a final model which performs well on both structure types, with an AUC-ROC 0.799 *±* 0.001 for predicted structures (Figure 2), and 0.807 *±* 0.001 when predicting solved structures (Supplementary Figure S1). We note that training separate models, namely using only solved or only predicted structures, does indeed improve performance slightly when tested on the same class (AUC-ROC 0.813 and 0.805 respectively), but comes at added complexity. To simplify comparison with other tools, we therefore chose the DiscoTope-3.0 version trained on both structure types for further analysis.

### 2.2 Benchmark comparison to state-of-the-art methods

To further test the effect of using predicted versus solved structures, we used the external test set of 24 antigens. These antigens share at most 20 % sequence similarity to both our own, BepiPred and ScanNet’s training datasets (see Methods). We benchmark against the structure-based tools ScanNet and SEMA, while including BepiPred-3.0, as a purely sequence-based and independent of the different structural variations, and a näıve predictor using relative surface accessibility as a score. We note that all benchmarked tools use the same definition for epitope residues, thus ensuring a fair comparison.

The precision and recall scores of the tools were calculated on this test set. The results of this evaluation are displayed in Figure 3. Here, DiscoTope-3.0 outperforms all other tools, for both predicted (AUC-PR 0.232 *±* 0.02 vs closest 0.177 *±* 0.02 BepiPred-3.0) and solved structures (0.223 *±* 0.02 vs closest 0.185 *±* 0.02 SEMA) (see Supplementary Figure S2 and Table 2 for more performance metrics). We note that DiscoTope-3.0 here strongly outperforms BepiPred-3.0, and point to the BepiPred-3.0 publication for an extensive benchmark demonstrating BepiPred-3.0 again outperforming epitope3D, Seppa-3.0, Ellipro and the previous version of DiscoTope-2.0.

**Fig. 3.**
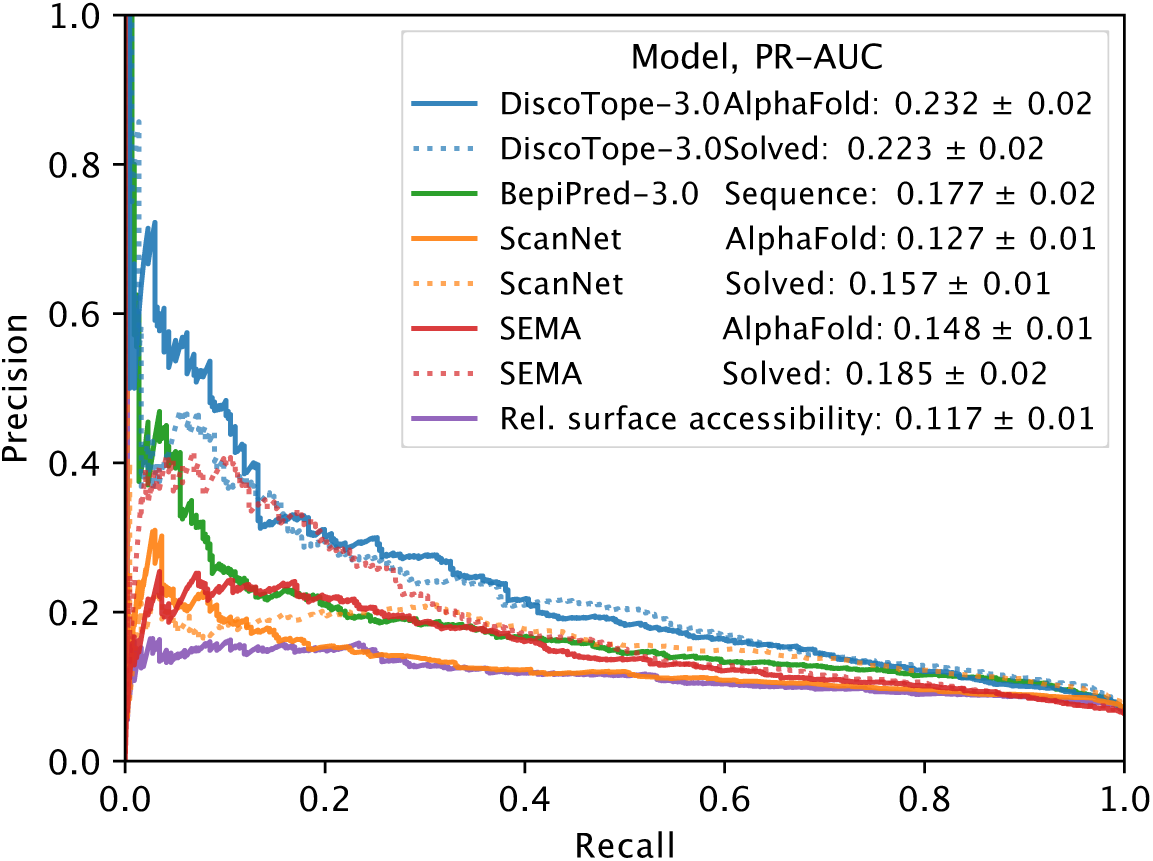
Improved performance on solved and predicted structures. AUC-PR curve plots on the external test set of 24 antigen chains, at most 20 % similar to the training set of all models. Structures provided as AlphaFold predicted, experimentally solved, or sequence in the case of BepiPred-3.0. Standard deviation calculated from bootstrapping 1000 times (see Methods). See Supplementary Figure S2 and Table 2 for additional performance metrics. Please see BepiPred-3.0 publication (Clifford et al., 2022) for its improved performance versus epitope3D, Seppa-3.0 and ElliPro.

**Table 1.** Feature overview. Input features for the XGBoost model architecture.

| Feature | Dimensions | Description |
| --- | --- | --- |
| ESM-IF1 embeddings | 512 | Inverse folding representations from input antigen structures |
| Antigen sequence | 20 | Amino-acid sequence, one-hot encoded |
| Antigen length | 1 | Length of sequence |
| AlphaFold quality score | 1 | Residue pLDDT score, as predicted by AlphaFold2 |
| Relative surface accessibility | 1 | Calculated by Shrake-Rupley algorithm |
| **Total:** | 535 | L x 535 |

**Table 2.**
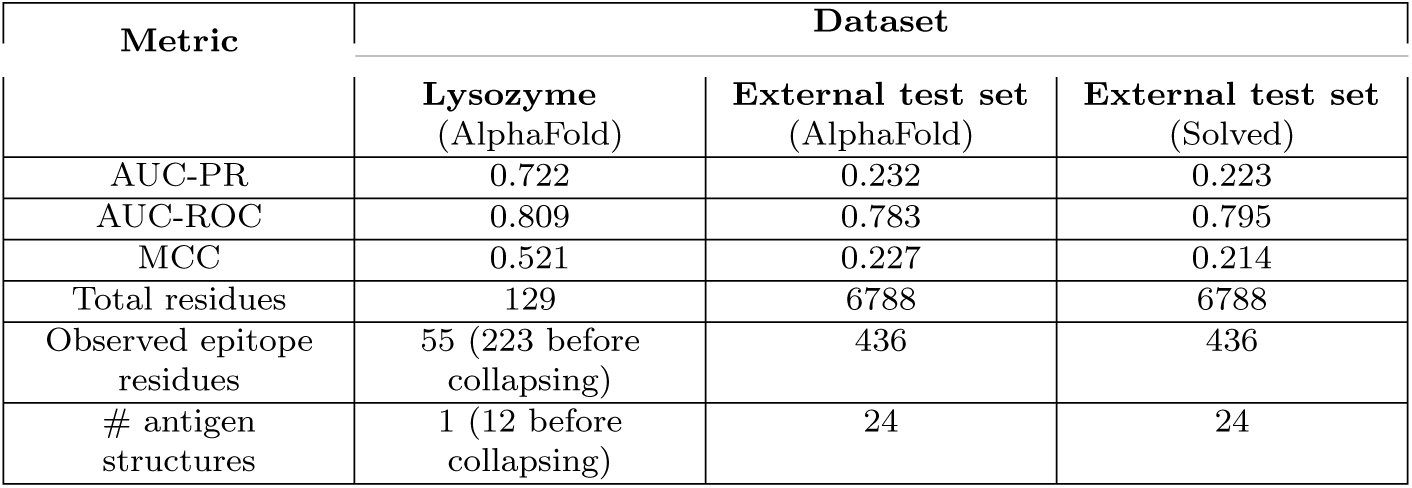
Performance on benchmarking datasets. Overview of external test set and lysozyme test sets for solved and AlphaFold predicted antigens. Matthew correlation coefficient (MCC) calculated at optimal sensitivity-specificity threshold using the Youden-index

We introduce a novel metric, the epitope rank score, primarily due to the need for a fair and normalized comparison among different tools, that operate with varying score scales. To put it simply, to calculate the epitope rank scores, we rank-normalize the scores for a given antigen, then find the mean rank for all observed epitopes in that antigen. For instance, a mean epitope rank score of 70% signifies that, on average, epitopes score in the upper 70th percentile of residue scores (see Methods). A typical real-case scenario for this metric, would be for users to submit individual antigens, and then to analyze the top scoring epitope residues, regardless of their specific scores. Using this metric, DiscoTope-3.0 consistently outperforms ScanNet, SEMA and BepiPred-3.0 in the case of both predicted and solved structures (Figure S3).

### 2.3 Robustness to relaxation and predicted structures

We note that DiscoTope-3.0’s performance is largely unaffected by the type of structures used for prediction. To further test the robustness of the tools to minor differences in the antigen structures, we performed an energy minimization on the solved structures using the software FoldX (Schymkowitz et al., 2005). This minimization only impacts the side chain, thus leaving the backbone of the native structure unaltered. The ESM-IF model does not use the side chain atoms in its structure representations, and consequently DiscoTope-3.0 should should not be affected by the relaxation process.

We observe that after side-chain relaxation in solved structures, ScanNet’s epitope rank scores are reduced by *∼*3.1 percentile points, while swapping solved for predicted structures leads to a loss of *∼*7.5 percentile points (see Methods). In contrast to this, DiscoTope-3.0 only loses *∼*0.1 and *∼*0.6 percentile points respectively, again indicating robustness to the modeling process (Supplementary Figure S4).

**Fig. 4.**
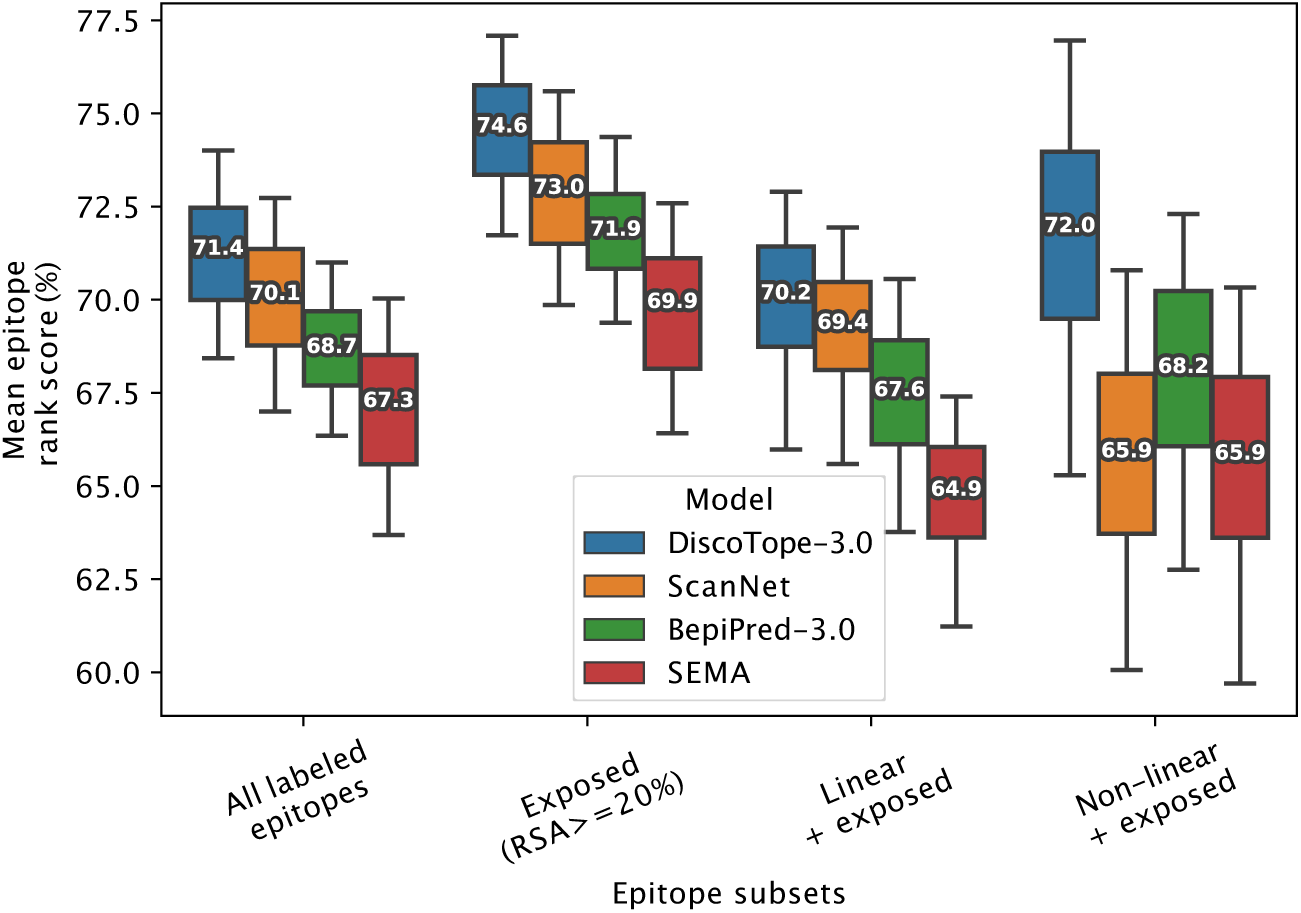
Improved performance on linear and non-linear epitopes. Mean epitope rank scores across antigens in the external test set, for the following epitope subsets: All labeled epitopes, Exposed (relative surface accessibility *<* 20%), Exposed Linear epitopes and exposed Non-linear epitopes (see text and Methods). Mean values calculated after bootstrapping 1000 times, with whiskers showing 95 % distribution range.

These observations can be attributed to the different ways the two models process structural features. ScanNet uses side-chain atomic coordinates explicitly, whereas DiscoTope-3.0 relies solely on the accuracy of backbone modeling. This difference suggests that some models, like ScanNet, might overfit to the specific orientations of side-chains present only in bound antibody-antigen complexes, information which would not be useful in predicting novel epitopes. By training models on both predicted and relaxed, solved structures, we can potentially avoid this overfitting and increase the generalizability of the models.

### 2.4 Improved prediction on exposed and non-linear epitopes

We also investigated if the structural information available to DiscoTope-3.0, ScanNet and SEMA affects the prediction of different types of epitopes. To this aim, epitopes were split into different sub-categories (Exposed, Buried, Linear and Non-linear). Exposed and Buried epitope residues are defined depending on whether their relative surface accessibility was above or below 20 %, respectively. Linear epitopes are defined as any group of 3 or more epitope residues found sequentially along the antigen sequence, allowing for a possible gap of up to 1 unlabeled residue in between. Finally non-linear epitopes were defined as epitopes not satisfying the conditions of the linear group.

The result of this performance evaluation in the external test set reveals improved performance of DiscoTope-3.0 across all epitope subsets (Figure 4). DiscoTope-3.0 performance is remarkably good for non-linear epitopes. In the case of buried epitopes (relative surface accessibility *<* 20 %), all models score poorly in the 30-37th percentile (not shown). This low performance is likely an artifact of the epitope labeling definition (shared between all tools), where inaccessible residues in proximity to the bound antibody are included in an epitope patch, despite not directly being involved in molecular interactions with the antibody.

### 2.5 Effect of predicted structural quality

Next, we investigate how the quality of the AlphaFold predicted structures affects the prediction of exposed epitopes. Overall, lower structural quality leads to small decrease in predictive performance (Figure 5), with high quality structures (pLDDT 95-100) having a mean epitope rank score of 84.2 %, and moderate quality structures (pLDDT 85-95) having a non-significant decrease in mean epitope rank score of 81.2 %. Only the group of antigens in the lowest quality pLDDT 60-85 group (*∼* 9 % of antigens) perform significantly worse, with a score of 75.5 % (*p <* 0.005). Fitting a linear model, the epitope rank score on average lowers by about 5 percentile points for every 10 point decrease in structural quality or pLDDT score (Figure 5B).

**Fig. 5.**
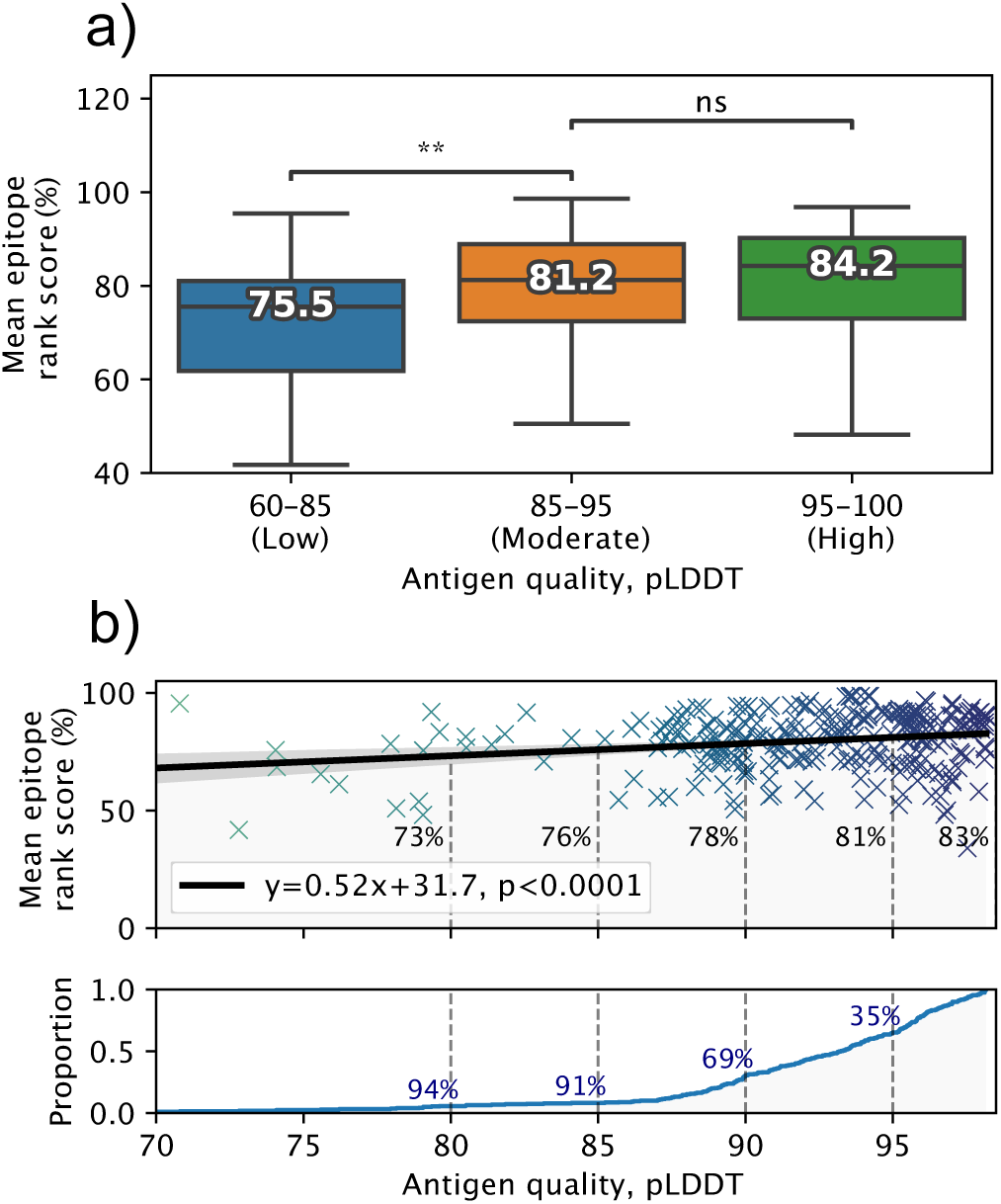
Effects of predicted structural quality. Validation set performance on AlphaFold predicted antigens dependent on predicted structural quality, excluding buried epitopes. a) Epitope rank score distribution for antigens split into increasing quality bins of mean antigen pLDDT 60-85, 85-95 and 95-100. Median value for each distribution is shown, with paired one-tailed t-test comparison (** = *p <* 0.005). b) Mean antigen pLDDT versus mean epitope rank score, with a fitted linear model shown in black. Below, cumulative distribution of mean antigen pLDDT, with a 91 % proportion exceeding a pLDDT of 85, and 35 % exceeding 95 respectively.

### 2.6 Calibrating scores for antigen length and surface area

We note that DiscoTope, BepiPred and SEMA exhibit a bias towards lower scores for longer antigen lengths (Pearson correlation -0.74, -0.71 and -0.51 respectively on external test set, not shown). If using a fixed threshold for binary epitope prediction, this results in most residues in shorter antigens being assigned as positives, while longer antigens may have all residues assigned as negatives.

To correct for this length bias, we calibrate antigen scores based on a predicted *µ* and standard deviation value, calculated from the antigen length and its mean surface score (see Methods and Supplementary figure S8). The calibrated scores demonstrate independence towards the antigen length, and clear separation of buried and exposed residues across antigens in the validation set S6. Furthermore, we find that calibrating the scores enables setting a fixed threshold that provides reliable epitope recall across shorter and longer antigens. For example, if we choose the 50th percentile calibrated score for exposed epitopes (in the validation set), and use this for binary epitope prediction on the lysozyme case study (see next section), we achieve an expected *∼* 50 % epitope recall (see Methods and Supplementary figure S7).

**Fig. 7.**
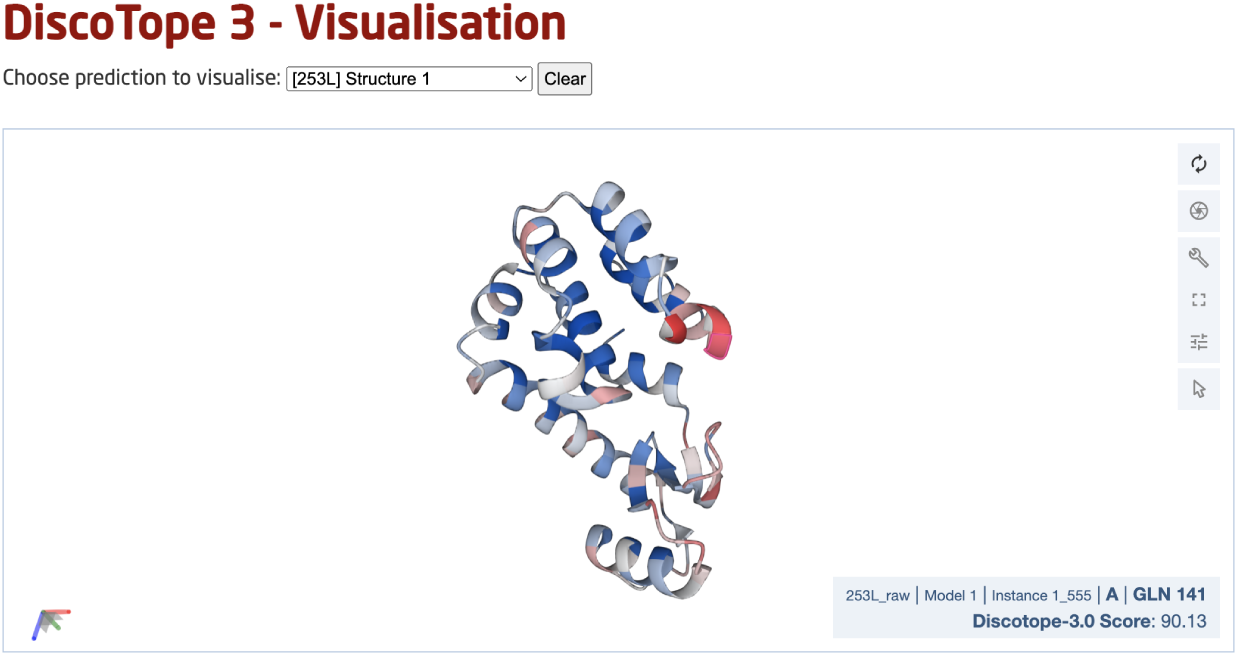
DiscoTope-3.0 web server interface. The web server provides an interactive 3D view for each predicted protein structure. DiscoTope-3.0 score on an example PDB, with increasing epitope propensity from blue to red. DiscoTope-3.0 is accessible at: https://services.healthtech.dtu.dk/service.php?DiscoTope-3.0

### 2.7 Lysozyme case study with collapsed epitopes

As a noteworthy test case, we evaluate the performance of DiscoTope-3.0 on lysozyme, a well-studied antigen extensively mapped against different antibodies. First, we identified 12 lysozyme chains with mapped epitopes at 90 % similarity to the chain C of the PDB structure 1A2Y. Next, DiscoTope-3.0 was re-trained excluding these chains. Next, we calculated an antibody hit rate, a ratio of on the number of times a given epitope residue was observed as an epitope across all of the 12 structures. Here, a score of 90 % means the same residue was observed as an epitope in 11 out of 12 of the chains, which is the case for 5 out of 129 residues.

Overall, we find that calibrated DiscoTope-3.0 scores correlate with the observed epitope count or antibody hit rate with a Spearman correlation of 0.58 (Figure 6). Fitting a linear model, we find that a 0.20 point increase in calibrated scores on average leads to a 10 % increase in the antibody hit rate (*p <* 0.0001, Supplementary Figure S5).

**Fig. 6.**
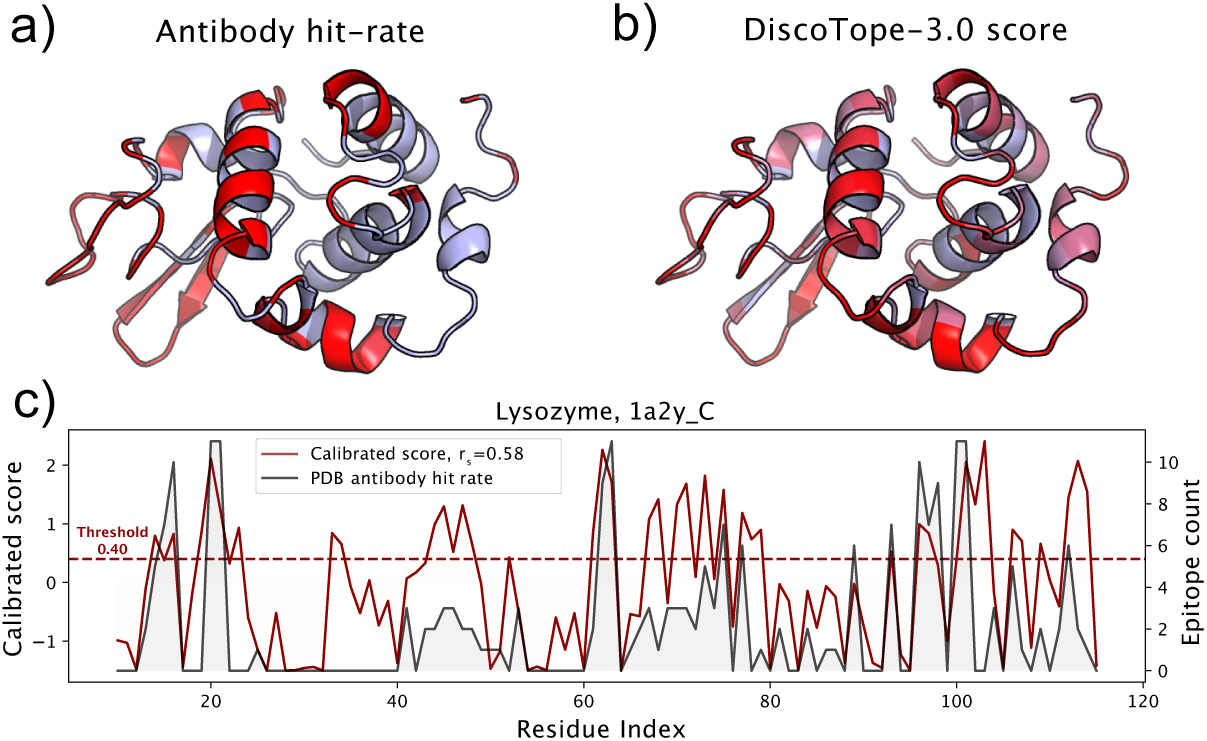
DiscoTope-3.0 score significantly correlates with antibody hit rate. Lysozyme epitope propensity as predicted by DiscoTope-3.0, excluding all lysozyme antigens from training. a) PDB antibody hit rate mapped AlphaFold predicted structure (chain 1a2y C), with increasing epitope propensity shown in red. b) DiscoTope-3.0 score c) Epitope propensity visualized across the antigen sequence. Calibrated DiscoTope-3.0 score and antibody hit rate (epitope counts) shown, as measured from aligning the 12 epitope mapped lysozyme sequences (Spearman R = 0.58). Additional performance metrics available in Supplementary table 2.

We note that the residues at positions *∼*30-40 (Figure 6) score highly in DiscoTope but lacked observed epitopes. Upon further investigation into the IEDB database, we found this region to be part of a discontinuous epitope patch (including K31, R32, G34, D36, G37, G40 . . . ) bound by a camelid antibody deposited under the PDB id 4I0C.

### 2.8 DiscoTope-3.0 web server

Finally, we deployed a DiscoTope-3.0 web server, which enables rapidly predicting epitopes on either solved or predicted structures. The web server currently accepts batches of up to 50 PDB files at a time, with any number of chains. Users may upload structures directly as PDB files, or automatically fetch existing structures submitted as a set of RCSB or AlphaFoldDB IDs. Output predictions are easily visualized through an interactive 3D view directly on the web server using Molstar (Sehnal et al., 2021), and predictions may be downloaded in both a CSV and PDB format.

## 3 Discussion

In this work, we present DiscoTope-3.0, a tool for improved B-cell epitope prediction. Our method exploits structure representations extracted from the ESM-IF1 inverse folding model. Extensive benchmarking of the tool demonstrated state of the art performance on both solved and predicted structures.

Importantly the performance, in contrast to earlier proposed structure-based models, was found to be maintained when shifting to predicted and relaxed structures. This observation is of critical importance since it removes the need for experimentally solved structures imposed in current structure-based models, and allows for predicted structures to be applied for accurate B-cell epitope predictions. This extends the applicability of the tool from the few thousand antigens for which a solved structure is available, to entire proteomes of available pre-computed structures modeled using AlphaFold-2.0.

We note that other structure-based tools perform worse than the sequence-based BepiPred-3.0 in cases where only predicted structures and their sequences are available. This may arise from sensitivity to the quality of predicted structures, or relying on signals only present in solved or unrelaxed structures. DiscoTope-3.0’s use of structure representations based on the protein backbone makes it robust to the predicted structural quality, and remarkably, able to perform similarly across solved, predicted and relaxed structures. It is, to the best of our knowledge, the first tool that presents highly accurate results on protein structural models.

We also find that other DiscoTope-like B-cell epitope prediction tools demonstrate a bias towards lower scores for longer antigens. After calibrating DiscoTope-3.0 scores for antigen length and surface residue scores, we provide calibrated score thresholds which provides the user with consistent expected epitope recall rates across shorter and longer antigens.

Finally, DiscoTope-3.0 interfaces with AlphaFoldDB and RCSB, enabling rapid batch processing across all currently cataloged proteins in UniProt and deposited solved structures. The web server is made freely available for academic use, accepting up to 50 input structures at a time, with any number of chains.

Our tool has been trained and evaluated on individual antigen chains. One could envision that, for multimeric antigen structures, it would be possible to further increase the tool performance by training and testing on the antigen complex. At this time, AlphaFold2 modeling accuracy for complexes is not yet on par with its accuracy on individual chains, and predicted complexes are not yet available in the AlphaFoldDB. As the science and technology behind the structural modeling progresses, it will be likely possible to further improve B-cell epitope predictions.

On the other hand, the positive-unlabelled learning strategy based on ensemble models and dataset bagging we use displays a remarkable boost in performance. We can imagine that, given the large dimension of the potential antibody space, the large gap between potential and observed epitopes will not be easily filled. An alternative strategy, that could circumvent this problem and provide valuable information to users, would be to perform antibody-specific epitope predictions. This approach has been tested by us and others in the past (Jespersen et al., 2019; Krawczyk et al., 2014), but the results are yet to provide a significant improvement in accuracy.

In summary, DiscoTope-3.0 is the first structure-based B-cell epitope prediction model that accepts and maintains state-of-the-art predictive power across solved, relaxed and predicted antigen structures. We believe this advance will serve as an important aid for the community in the quest for novel rational methods for the design of novel immunotherapeutics.

## 4 Data and code availability

DiscoTope-3.0 web server, downloadable package and training datasets are freely available for academic use.

- Web server DTU: https://services.healthtech.dtu.dk/service.php?DiscoTope-3.0
- Web server Biolib: https://biolib.com/DTU/DiscoTope-3/
- Code availability: https://github.com/Magnushhoie/discotope3web

## 6 Author contributions

MH, PM, MN contributed to conception and design of the study. MH, FG, JJ, CW implemented the methodology and software. FG, MH implemented the web server. MH, JJ, CW performed the statistical analysis and visualization of results. MH, MN, PM wrote sections of the manuscript. PM, MN, OW provided supervision. All authors contributed to manuscript revision, read, and approved the submitted version.

## 8 Methods

### 8.1 Training and evaluation of DiscoTope-3.0

The antigen training dataset as presented in BepiPred-3.0 was used as the starting point for our work. The dataset consists of 582 AbAg crystal structures from the PDB, filtered for a minimum resolution of 3.0 Å and R-factor 0.3. Epitopes are defined as any antigen residue containing at least 1 heavy atom within 4 Å of an antibody heavy atom. From this dataset, using the tool MMseqs2, we first remove any sequences with more than 20 % sequence identity to the BepiPred-3.0 test set, resulting in 1406 chains. Next, the antigen sequences are clustered at 50 % sequence identity. Each cluster has then been selected to be part of the validation (281 chains) or the training set (1125 chains).

In the ablation study, single XGBoost models (Chen and Guestrin, 2016) with default parameters were trained using representations from either the predicted structure or antigen sequence respectively. When testing feature combinations, ensemble size and effect of training on solved and predicted structures, error bars were estimated from re-training 20 times.

Hyperparameters for the XGBoost models were manually optimized using the validation dataset. We used the final parameters n estimators=200, max depth=4, learning rate=0.3 and subsample=0.50. The gpu hist tree method was used for faster training on a GPU.

### 8.2 Dataset bagging and ensemble training

When sampling residues for each model in the ensemble, we randomly select 70 % of available observed epitopes (positives) across the training dataset, then sample unlabelled residues (negatives) with a ratio of 5:2. When using both predicted and solved structures, these were sampled at a 1:1 ratio.

Ensembles were constructed by iteratively training independently trained XGBoost models on the randomly sampled datasets. When training an ensemble, we set a different random seed each time.

### 8.3 ESM-IF1 and ESM-2 representations

To generate per-residue ESM-IF1 structure representations, antigen structures were first split into single chains, and these inputted into ESM-IF1 following the instructions as listed on the official repository (Research, Research).

1. `import esm.inverse_folding`
2. `structure = esm.inverse_folding.util.load_structure(fpath, ↪ chain_id)`
3. `coords, seq = ↪ esm.inverse_folding.util.extract_coords_from_structure(structure)`
4. `rep = esm.inverse_folding.util.get_encoder_output(model, ↪ alphabet, coords)`

For per-residue ESM-2 sequence representations, sequences were first extracted from all antigen chains and stored in a FASTA format. Next, the FASTA file was provided as input to the official extract.py script (Research, Research) using the pre-trained ESM-2 650M parameter model.

`./extract.py --model_location esm2_t33_650M_UR50D \`

`--fasta_file sequences.fasta \`

`--include per_tok \`

`--output_dir output/`

### 8.4 Feature calculation and data filtering

Each isolated chain was processed as a single PDB file with ESM-IF1, extracting for each residue its latent representation from the ESM-IF1 encoder output. pLDDT values were either extracted from the PDB files in the case of AlphaFold structures, or set to 100 for solved structures. In the case of training on both solved and predicted structures, we include a binary input feature set to 1 if the input is an AlphaFold2 model, and 0 for solved structures.

Residue solvent accessible surface area was calculated using the Shrake-Rupley algorithm using Biotite (Kunzmann and Hamacher, 2018), with default settings, and converted to relative surface accessibility using the Sander and Rost 1994 (Rost and Sander, 1994) scale as available in Biopython (Cock et al., 2009).

When training DiscoTope-3.0, we removed any antigen with less than 5 or more than 75 epitope residues, as well as PDBs with a mean pLDDT score below 85 or residues with a pLDDT below 70. No data filtering was performed during evaluation on the validation and external test datasets.

### 8.5 Calibration of DiscoTope-3.0 scores

When using calibrated scores, each antigen’s DiscoTope-3.0 scores are normalized using the following formula:

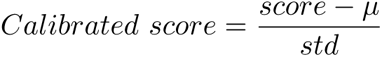

The values for *µ* and std are calculated for each antigen, using two separate linear generative additive models (GAMs) (Servén D., 2018) fitted on the validation set. The length to µ model is fitted on antigen length versus mean score of antigen surface residues (RSA *>* 20 %), while the surface mean to std model is fitted on antigen mean surface residue score versus standard deviation of the same scores (Supplementary Figure S8).

### 8.6 External test set generation and evaluation

The external test set, used for comparing our tool to ScanNet and BepiPred-3.0, consists of solved antibody-antigen complexes deposited in either SAbDab and the PDB after April 2021 (collection date June 2022). Any antigen with more than 20% sequence identity to the training datasets used in this work, in ScanNet, or in BepiPred-3.0 were removed using MMseqs2. We annotated epitopes using the same approach as in BepiPred-3.0, which is common to all the tools.

We submitted either solved or AlphaFold2 predicted structures to the Scan-Net web server (Tubiana et al., 2022a), using the antibody-antigen binding mode and otherwise default parameters. BepiPred-3.0 predictions were generated from its online web server (Clifford et al., 2022) using the antigen sequence and default parameters.

When evaluating DiscoTope-3.0 on the external test set, we retrained the final model with an ensemble size of 100, on the full training and

### 8.7 AlphaFold2 modeling and structural relaxation

Sequences for each antigen chain containing at least 1 epitope were extracted and modeled with the ColabFold implementation of AlphaFold2 at default settings. We picked the top ranking PDB after 5 independent iterations of 3 recycles, as ranked by AlphaFold2’s internal quality measure.

For relaxation of the solved structures we used the foldx 20221231 version of FoldX, with the RepairPDB command for relaxing residues with bad torsion angles, van der Waals clashes or high total energy.

### 8.8 Data analysis

To calculate the mean epitope rank score, the predicted residue scores for an antigen were first ranked in ascending order. Next, we calculated the average of the rank scores for all epitope residues.

Exposed epitopes were defined as all epitopes with a relative surface accessibility exceeding 20 %, while the remaining epitopes were defined as buried.

When reported, significance testing was performed with a one-sided paired t-test using scipy.stats.ttest rel (Virtanen et al., 2020). The linear model on the mean antigen pLDDT vs mean epitope rank scores was fitted using a linear least-squares regression model (scipy.stats.linregress) with two-sided alternative hypothesis testing.

For confidence estimation with bootstrapping, the dataset was sampled fully with replacement 1000 times, with the bootstrapped datasets used to calculate means, epitope rank scores and standard deviation values.

## Supplementary Figures

**Fig. S1.**
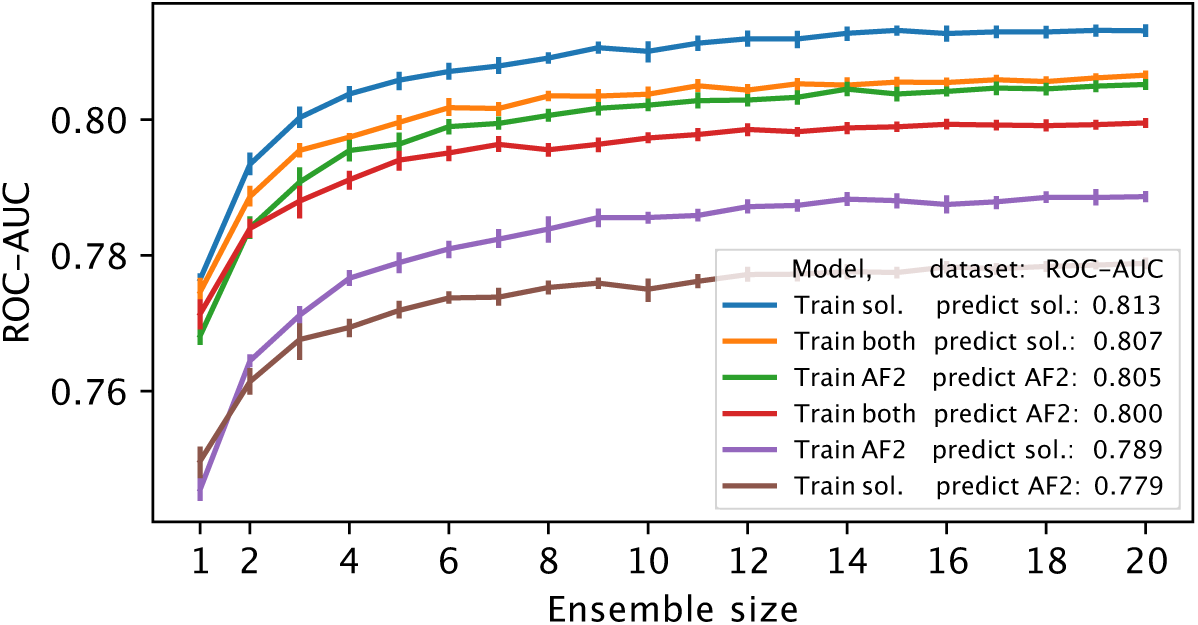
Effect of ensemble size. Validation set gain in AUC-ROC from ensembling the full-feature model. Performance graphs are shown for training on either experimentally solved, AlphaFold predicted or both structures, and then evaluated on either the solved or predicted structure validation set.

**Fig. S2.**
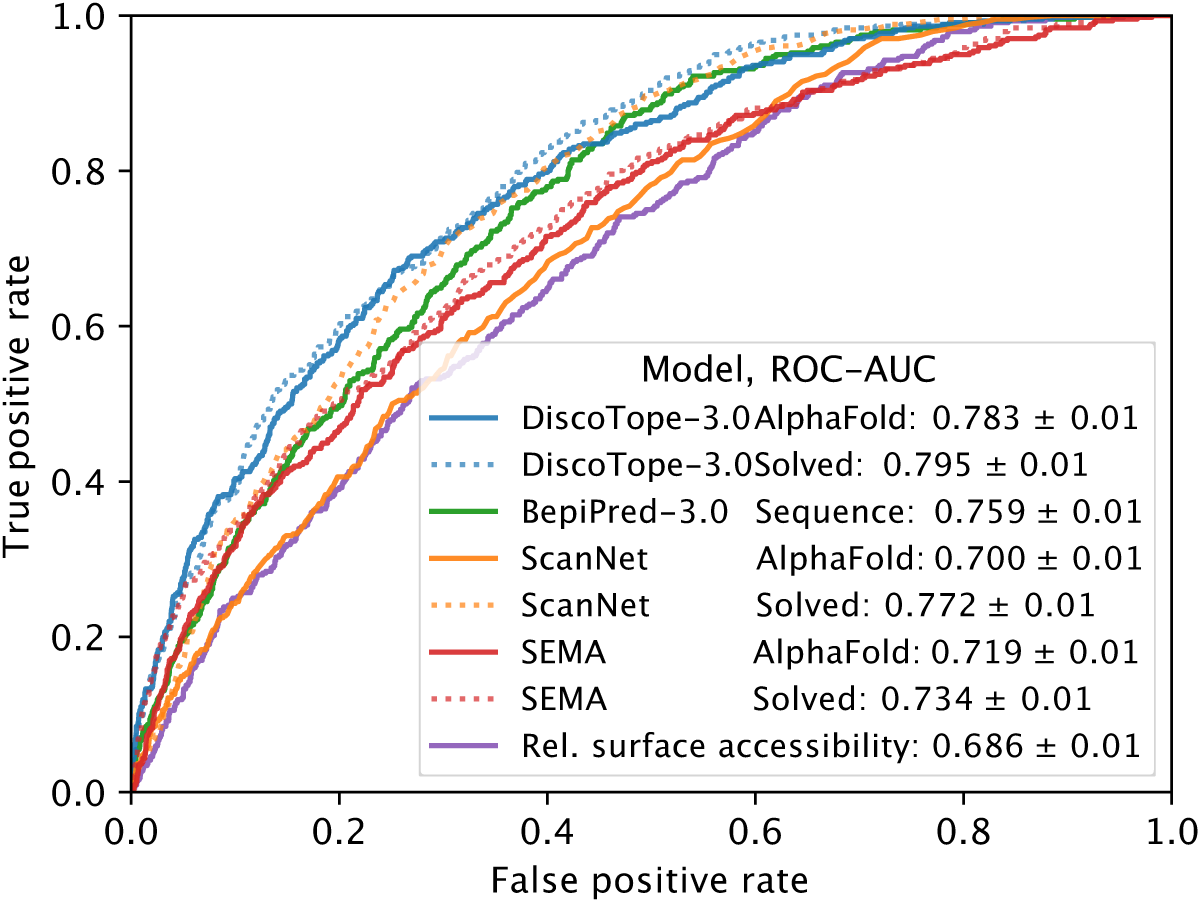
External test set AUC-ROC. Test set AUC-ROC, as evaluated on 24 antigens modeled with AlphaFold. For PR-AUC see Figure 3.

**Fig. S3.**
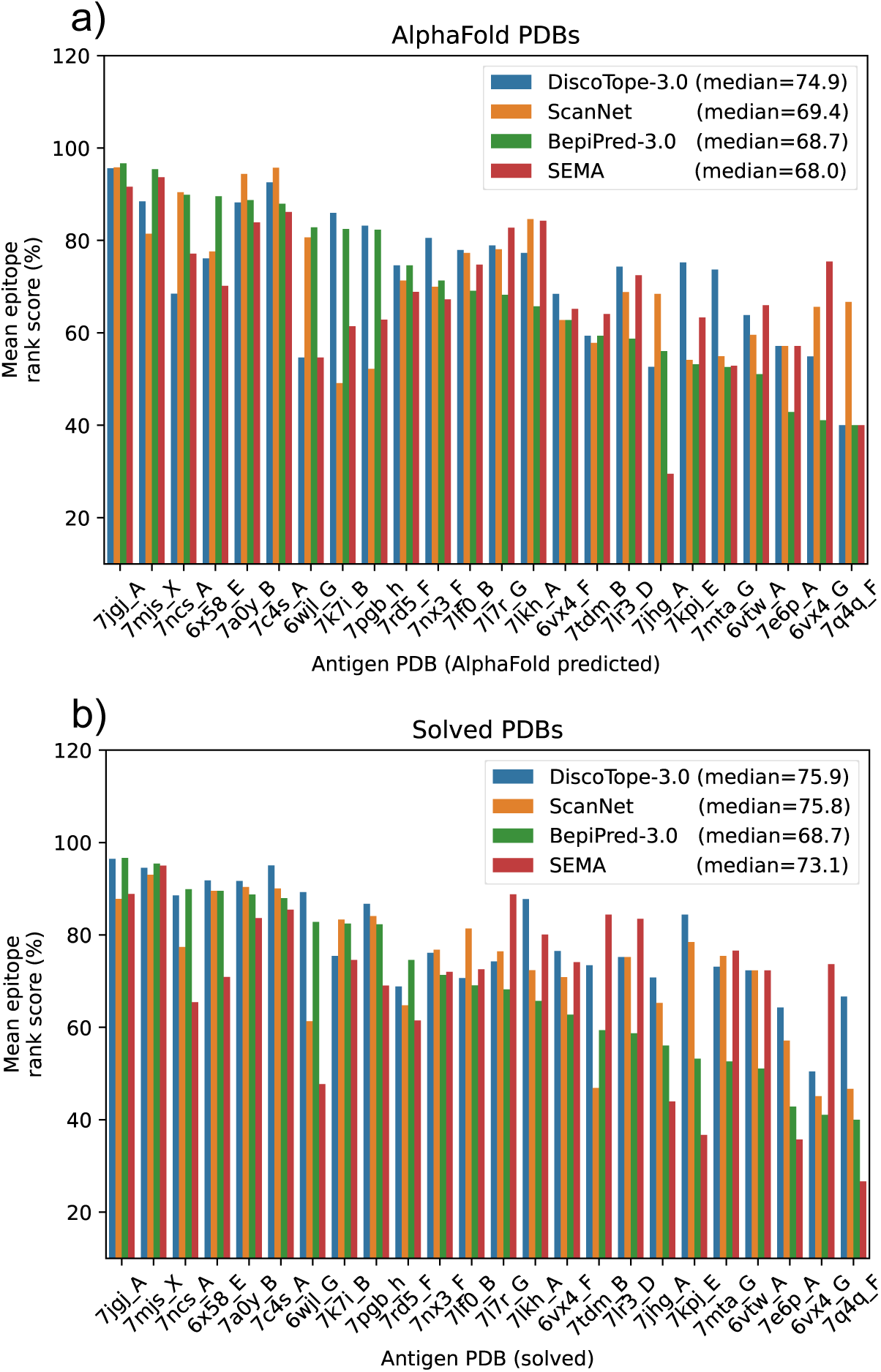
External test set PDB performances. Evaluation on 24 antigens modeled with AlphaFold (left) or experimentally solved structures (right). BepiPred-3.0 performances on antigen sequences only.

**Fig. S4.**
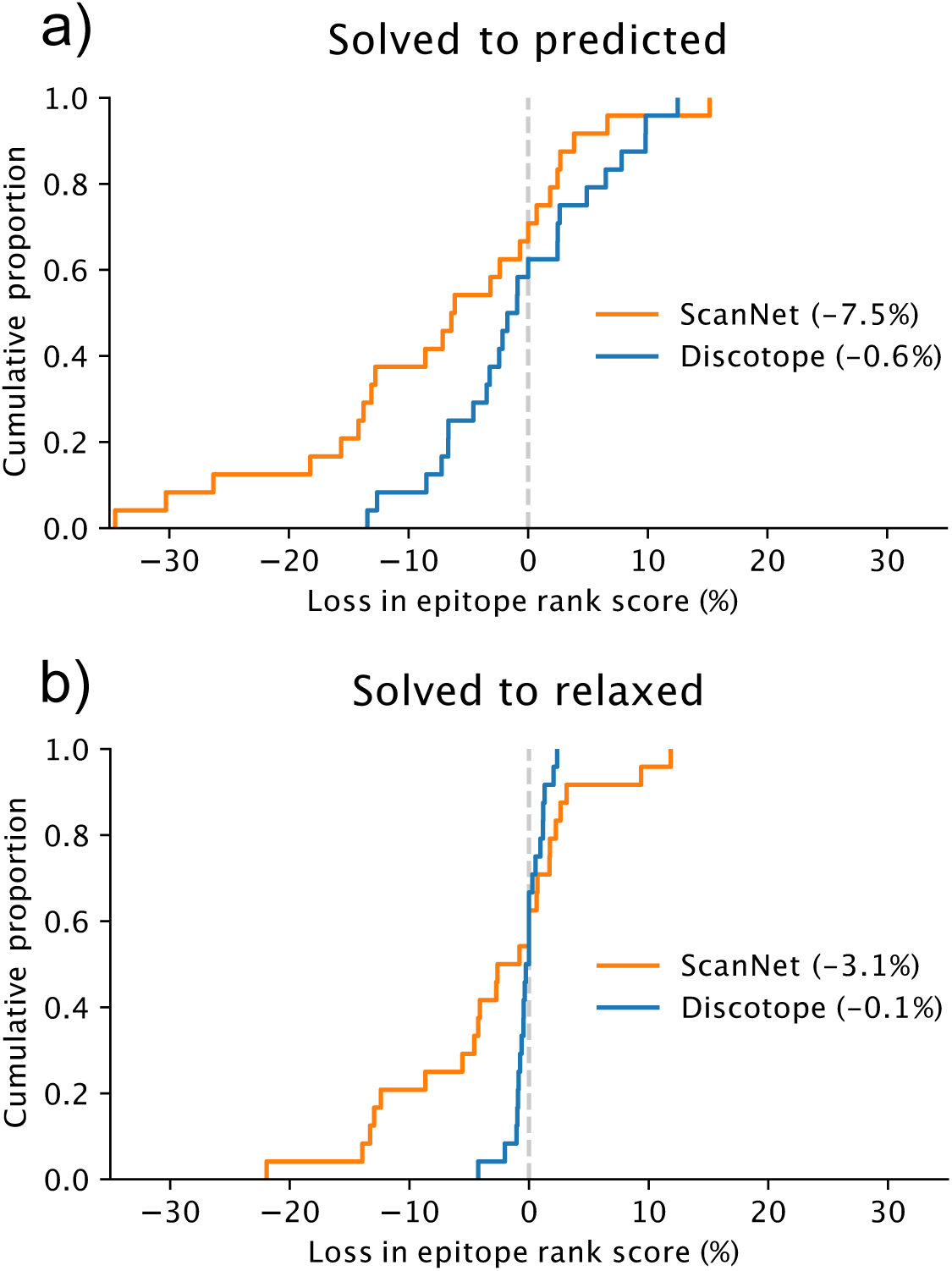
DiscoTope-3.0 is robust towards modeling and relaxation. External test set change in mean epitope rank scores across PDBs, when (a) swapping predicted structures with their original solved structure or (b) solved structures with the same structure after FoldX relaxation (see Methods). Mean performance loss shown in percent.

**Fig. S5.**
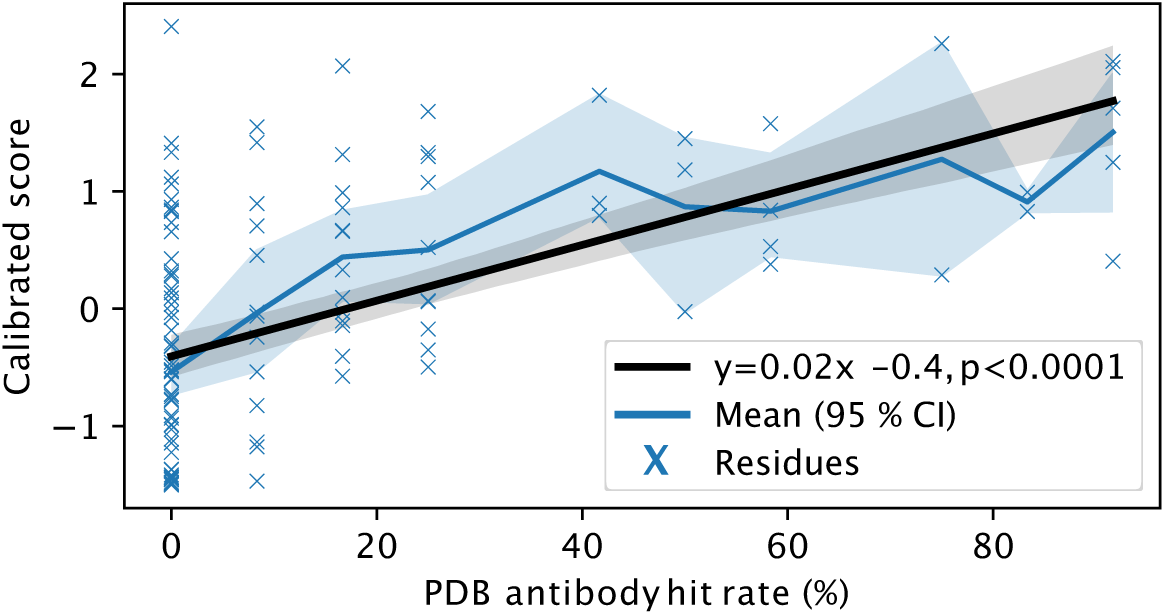
DiscoTope-3.0 score significantly correlates with antibody hit rate. Lysozyme case study on 1a2y C, showing PDB antibody hit rate (ratio of times an epitope residue is observed across all 12 lysozyme chains) versus calibrated DiscoTope-3.0 scores. Model is trained excluding all lysozyme structures from training (see Methods).

**Fig. S6.**
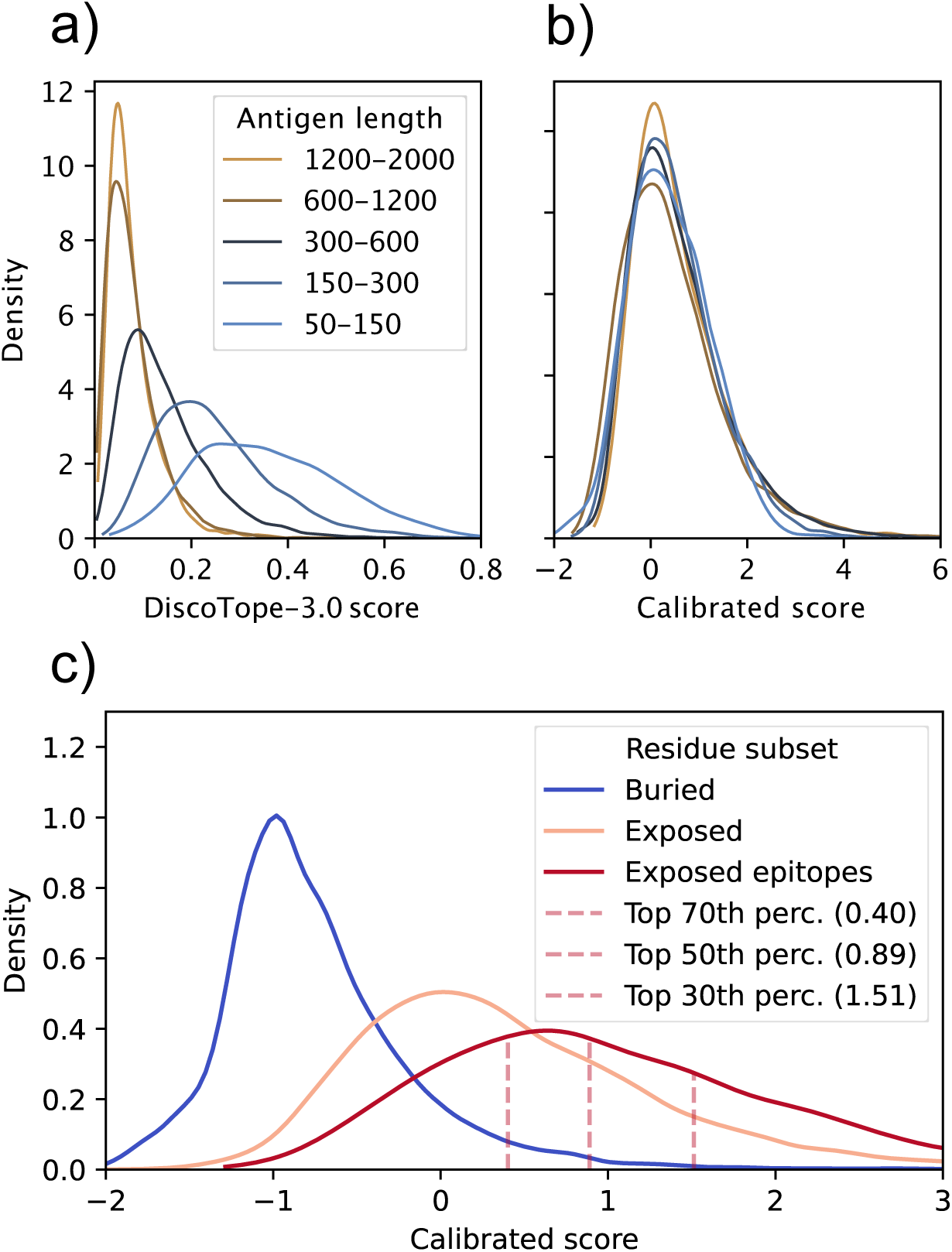
DiscoTope-3.0 calibrates for antigen length and surface area. Uncalibrated DiscoTope-3.0 surface scores are biased towards the antigen length. A) Validations set DiscoTope-3.0 score distributions before normalization and B) after correcting for antigen length and surface scores (see Methods). C) Calibrated score distributions in the validation set, for buried residues, exposed residues (relative surface accessibility *>* 20 %) and exposed epitopes. The top 70th, 50th and 30th percentile scores for exposed epitopes are shown in red dashed lines (A and B), as suggestive thresholds for binary epitope prediction.

**Fig. S7.**
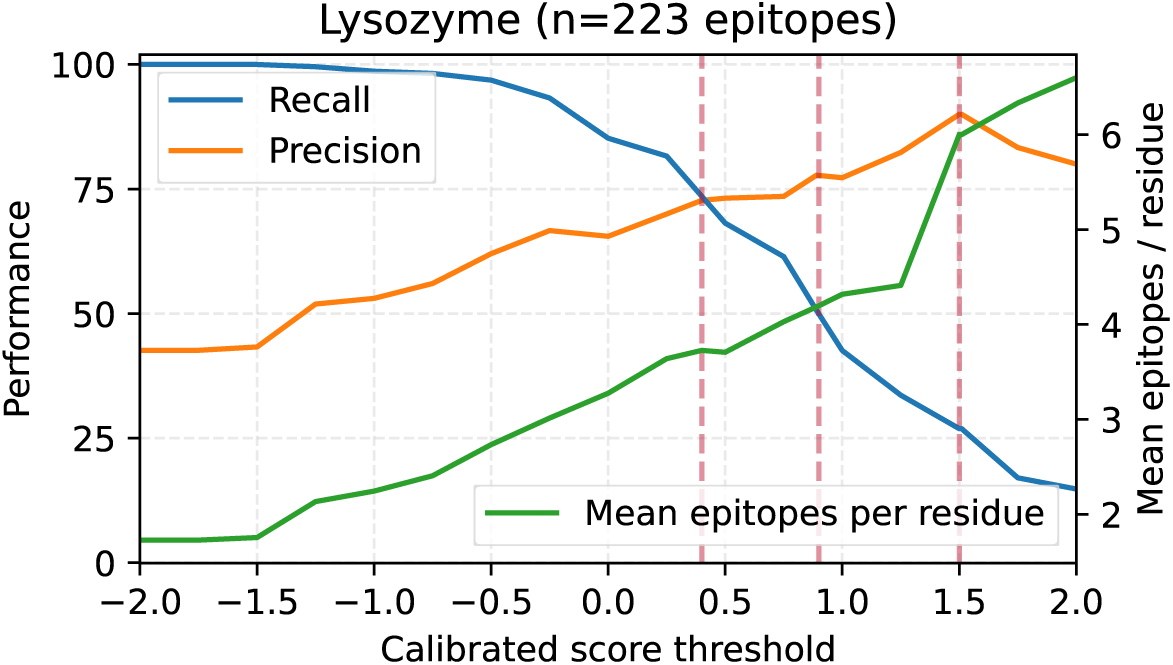
Benchmarking calibrated score thresholds on Lysozyme. Binary epitope prediction performance on the collapsed lyzosyme dataset, for different calibrated score thresholds. Recall of total observed epitopes shown in blue, with precision for any epitopes in Denmark range. Green line shows the median epitope count per residue for residues above the given threshold (maximum 12). Red lines shown for the previously mentioned top 70th, 50th and 30th exposed epitope percentile scores from the validation set (Fig S6).

**Fig. S8.**
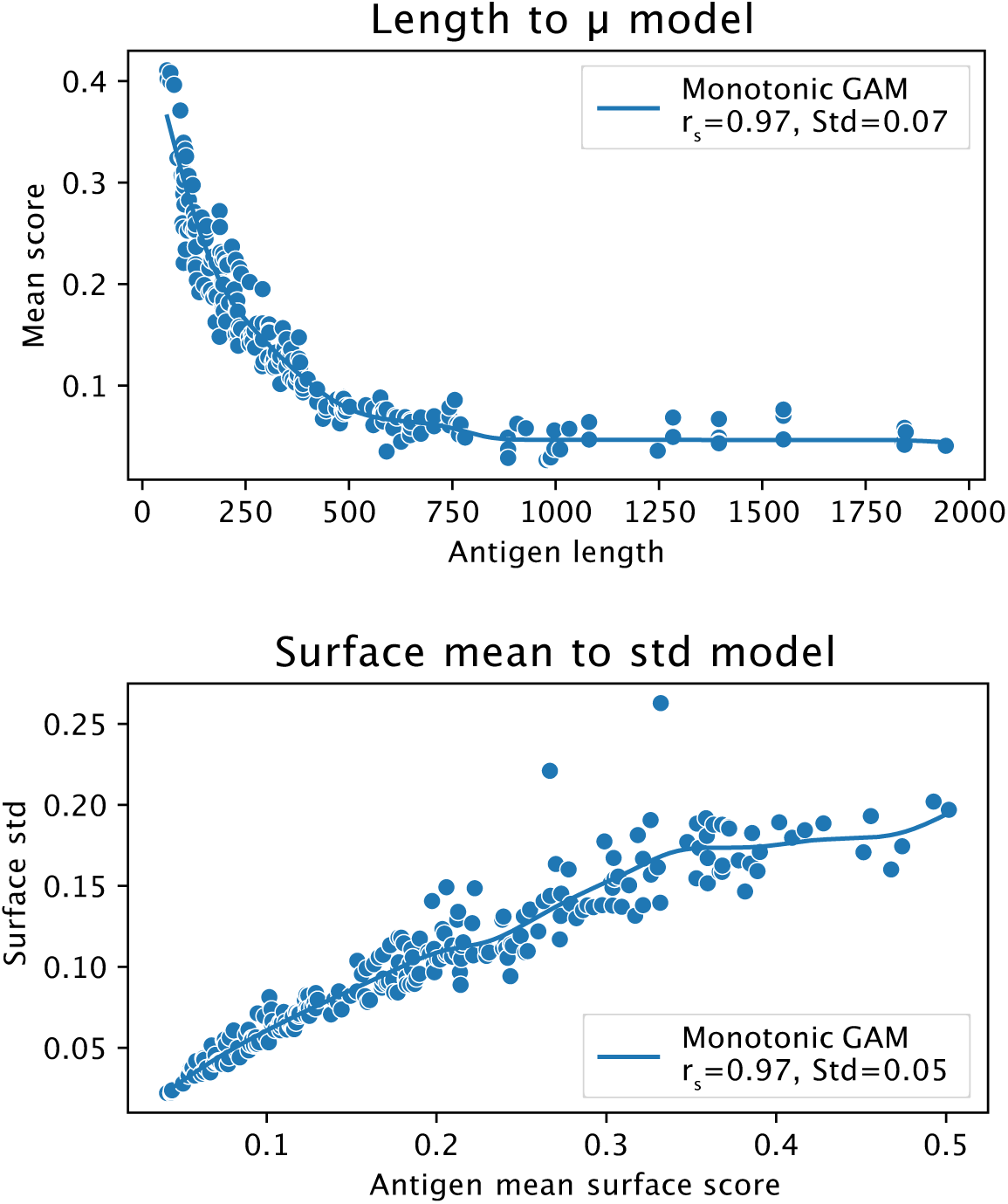
Fitted GAM models for calibrating scores. Length to *µ* and surface to std fitted GAM models on the validation set, used for calibrating DiscoTope-3.0 scores (see Methods).

